# Mutational load and the functional fraction of the human genome

**DOI:** 10.1101/785865

**Authors:** Benjamin Galeota-Sprung, Paul Sniegowski, Warren Ewens

## Abstract

The fraction of the human genome that is functional is a question of both evolutionary and practical importance. Studies of sequence divergence have suggested that the functional fraction of the human genome is likely to be no more than ∼15%. In contrast, the ENCODE project, a systematic effort to map regions of transcription, transcription factor association, chromatin structure, and histone modification, assigned function to 80% of the human genome. In this paper we examine whether and how an analysis based on mutational load might set a limit on the functional fraction. In order to do so, we characterize the distribution of fitness of a large, finite, diploid population at mutation-selection equilibrium. In particular, if mean fitness is ∼1, the fitness of the fittest individual likely to occur cannot be unreasonably high. We find that at equilibrium, the distribution of log fitness has variance *nus*, where *u* is the per-base deleterious mutation rate, *n* is the number of functional sites (and hence incorporates the functional fraction *f*), and *s* is the selection coefficient of deleterious mutations. In a large (*N* = 10^9^) reproducing population, the fitness of the fittest individual likely to exist is 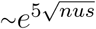. These results apply to both additive and recessive fitness schemes. Our approach is different from previous work that compared mean fitness at mutation-selection equilibrium to the fitness of an individual who has no deleterious mutations; we show that such an individual is exceedingly unlikely to exist. We find that the functional fraction is not very likely to be limited substantially by mutational load, and that any such limit, if it exists, depends strongly on the selection coefficients of new deleterious mutations.

## 1 Introduction

The total proportion of the human genome that is functional has been a question of intense interest. It has long been known that, across different species, genome size does not bear a close relationship to apparent complexity: for example, the lungfish genome is 60 times larger than the human genome, and there is a three-order-of-magnitude range of genome sizes in angiosperms. This is the C-value paradox (Thomas 1971; reviewed in Gregory 2005).

A natural definition of “functional” is “selected for at the organismal level”, which implies the possibility of deleterious mutation (Graur 2013). Evolutionary studies that examine divergence from related organisms (reviewed in Ponting and Hardison 2011, and see Rands *et al.* 2014), some of which also utilize intraspecies variation (*e.g.* Gulko *et al.* 2015, Ward and Kellis 2012), suggest that 3-15% of the human genome is subject to purifying selection, with methods that account for rapidly evolving yet still constrained sequences (e.g. Meader *et al.* 2010) tending to fall on the higher end of this range.

The ENCODE project (Dunham *et al.* (ENCODE) 2012) was a large-scale systematic effort to map regions of transcription, transcription factor association, chromatin structure and histone modification in the human genome. Regions assigned to any of these mappings were considered by the ENCODE authors to be functional, leading to a total estimate of 80% of the human genome as functional. The discordance between the ENCODE estimate and those of other studies, together with ENCODE’s expansive definition of functionality—one seemingly divorced from an evolutionary approach—led to criticism (Doolittle 2013; Graur *et al.* 2013) of the ENCODE estimate. Indeed, such a high fraction of functionality would be difficult to reconcile with the fact that one half to two thirds of the human genome consists of inactivated transposable elements (International Human Genome Sequencing Consortium 2001, de Koning *et al.* 2011). Nor does a high estimate for the proportion of the human genome that is functional help to resolve the C-value paradox (Doolittle 2013).

Consideration of mutational load may set a limit on the functional fraction. By comparing the population mean fitness at mutation-selection equilibrium to that of an individual who possesses no deleterious mutations, Graur (2017) reached the conclusion that, for likely values of the human per-base deleterious mutation rate, the functional fraction must be small.

In this paper, we present a different approach to analyzing mutational load and the human functional fraction. We do not take the fitness of an individual with zero deleterious mutations to be a meaningful value, because in a finite population of realistic size such an individual will never exist. Instead, we consider the fitness of the fittest individual likely to exist in a finite population. We conclude—while making no claims about the actual functional fraction as determined by comparative studies—that a mutational load argument is unlikely to set a low limit on the functional fraction of the human genome, and that any attempt to set such a limit must take into account the fitness effects of new deleterious mutations.

## 2 Theoretical development

### 2.1 The mutational load

There are various definitions of a genetic load “*L*” in the literature. Perhaps the most frequently used definition (Crow 1970) is

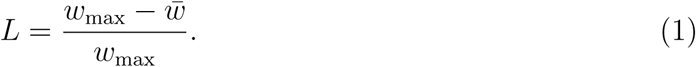

Here 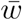 is the mean population fitness and *w*_max_ is the fitness of individuals of the fittest possible genotype. There are many kinds of loads; in this case we are concerned with the load due to recurrent deleterious mutation, *i.e.* the mutational load. Note that the form of (1) leaves open the question of what exactly is represented by *w*_*max*_.

Although (1) is a useful measure of the proportion by which mean fitness is lower than the maximum, we find that the key quantity to consider in relation to the functional fraction of the genome is

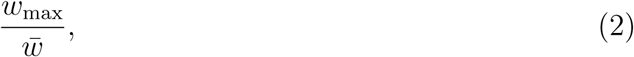

the factor by which the maximum fitness is greater than the mean. When mean fitness is 1, this reduces simply to *w*_*max*_. In our analysis, we assign *w*_*max*_ to the fitness of the fittest individual likely to exist in a finite population. The analysis of Graur (2017) is rather different, but is concerned with the quantity 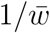 where *w*_*max*_ = 1; hence both analyses are ultimately concerned with the value of (2).

### 2.2 Fitness model, assumptions, and parameter values

We define a functional site as one at which a deleterious mutation is possible, and these are the only sites that we consider. For a single functional site, we define *A* as the normal nucleotide at this site and *M* as the mutant nucleotide that imparts a reduced fitness when homozygous: relative to the fitness of the *AA* genotype, the fitness of the *MM* genotype is multiplied by 1 − *s*, where *s* > 0. The fitness of the heterozygote depends on whether an additive or recessive fitness arrangement is employed. We assume a mutation rate of *u* from *A* to *M*.

The value of *u* is very small and following common practice we ignore terms of order *u*^2^ in our development, as well as the exceedingly small effect of back mutations from *M* to *A*. Empirical estimates of human mutation rates include all mutations that occur. However, not all mutations, even at a functional site, are necessarily deleterious. We use *v* to denote the empirically estimated rate of mutation per base pair in the human genome and *p* to denote the probability that a mutation is deleterious. Then *u* = *vp* is the probability of a deleterious mutation at a functional site. We adopt the value *v* = 1.2 × 10^−8^ in agreement with recent studies (Lesecque *et al.* 2012; Besenbacher *et al.* 2015; Milholland *et al.* 2017). Graur (2017) provides evidence that 0.4 is a reasonable value for *p*, the probability that a mutation at any functional site is deleterious, and we also adopt this value.

In addition to the parameters *v, u, p* and *s* defined above, we define *g* as the number of base pairs in the diploid human genome and *f* as the proportion of sites in the genome that are functional. This implies that the number of functional diploid sites in the genome is 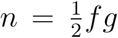, with the factor 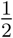 accounting for the fact that a site encompasses two homologous base pairs. We consider only these *n* functional sites. We adopt the recent estimate *g* = 6.4 × 10^9^ (Schneider *et al.* 2017), except when directly comparing to Graur (2017) in which a slightly different value is used.

It is harder to choose a value for *s*, first because published estimates of *s* rely on indirect methods with large resulting standard errors, and second because the value of *s* surely differs from site to site, and probably substantially. Lesecque *et al.* (2012) recognize the site-to-site variation in *s* and illustrate their analysis by considering the cases *s* = 10^−3^, *s* = 10^−2^ and *s* = 10^−1^. This corresponds to the Eyre-Walker *et al.* (2006) and Eyre-Walker and Keightley (2007) estimates that in humans most functional sites have a value of *s* in the range 10^−3^ to 10^−1^. Boyko *et al* (2008) find that roughly one third of nonsynonymous mutations have *s* < 0.0001, one third have 0.00001 < *s* < 0.01, and one third have *s* > 0.01. For convenience, we choose a value *s* = 10^−2^ for some of our calculations; but we report results for a range of values from *s* = 10^−2^ to *s* = 10^−4^ (Table 1).

**Table 1:**
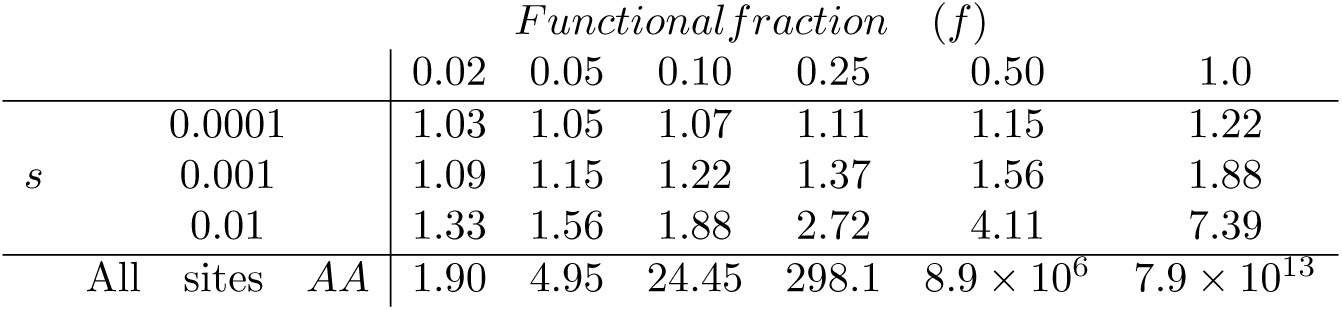
Fitness values *w*_*max*_ of the fittest individual likely to occur in a population of size *N* = 10^9^, according to (12), for a range of values for the functional fraction and the selection coefficient, assuming that mean fitness is standardized to 1. The parameters *u* and *g* are as given in (3). The bottom row shows, for comparison, the fitness of the fittest possible individual in the additive model (13), who is *AA* at all sites—but who will never exist in a real finite population.

| | | | *Functional fraction* ($f$) | | | | | |
| --- | --- | --- | --- | --- | --- | --- | --- | --- |
|  |  |  | 0.02 | 0.05 | 0.10 | 0.25 | 0.50 | 1.0 |
| $s$ | 0.0001 | | 1.03 | 1.05 | 1.07 | 1.11 | 1.15 | 1.22 |
|  | 0.001 |  | 1.09 | 1.15 | 1.22 | 1.37 | 1.56 | 1.88 |
|  | 0.01 |  | 1.33 | 1.56 | 1.88 | 2.72 | 4.11 | 7.39 |
| All | sites | $AA$ | 1.90 | 4.95 | 24.45 | 298.1 | $8.9 \times 10^6$ | $7.9 \times 10^{13}$ |

We will later assume a reproducing population size of *N* = 10^9^. This is a value that the human population has only recently attained, and is very conservative in the sense that it is the one least favorable to our arguments that the functional fraction is not very limited by mutational load. To summarize, unless noted otherwise we will use the following parameter values in some of our calculations:

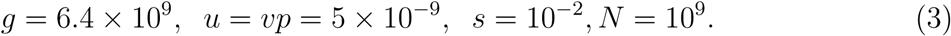

The quantity *nus* will enter often into our calculations below. For example, with the parameter values in (3) and with *f* = 0.05, a value included in Table 1, *nus* = 0.008.

The features of our model as defined above are compatible with the model of Graur (2017), and we have accordingly adopted much of the same notation, except that for notational convenience we use *u* where he uses *µ*_*del*_.

## 3 Results

### 3.1 The additive case

We first consider the additive fitness, or no dominance, case, since other authors appear to focus on this case. For any given functional site the relative, or proportional, genotype fitnesses are of the following additive form:

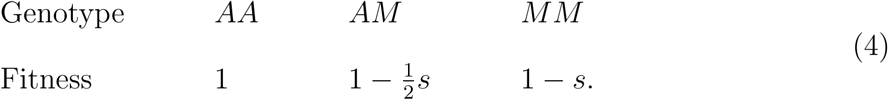

The equilibrium frequency of *A* at each such site is 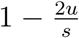 so that population mean fitness at each site is the well-known value 1 − 2*u* (Crow and Kimura 1970).

The size of the human population has expanded greatly in the last 10,000-100,000 years, but this is a relatively recent phenomenon on evolutionary time scales. It is reasonable to assume that over the long time scales in which the fundamental features of the genome have been shaped, mean absolute fitness has been close to 1, *i.e* the population size has been constant. Other authors (e.g. Haldane 1937; Graur 2017) have made this assumption, which we follow by multiplying the fitnesses by a common factor such that the mean fitness is 1. This normalization leads to the following well-known genotype fitnesses and frequencies (Crow and Kimura 1970):

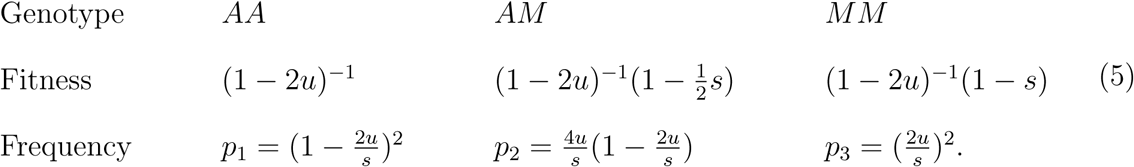

According to (5), the fitness of an individual who is *AA* at all *n* functional sites is (1−2*u*)^−*n*^ ≈ *e*^2*nu*^. A main point of this paper is that no individual of this genotype will ever exist in a natural population, so no calculation that depends upon such an individual (i.e. assigns *w*_*max*_ to such a value) is empirically relevant. The probability that a randomly chosen individual is *AA* at all *n* functional sites is 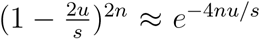. With the parameter values in (3) and with *f* = 0.05 this is about *e*^−320^. Similar calculations arise with other plausible parameter values. No individual who is *AA* at anything approaching *n* sites will ever appear in a natural population.

#### 3.1.1 The distribution of whole-genome fitnesses

With these considerations in mind, our aim is to calculate the whole-genome distribution of fitnesses and establish a methodology for defining an upper limit to *f* based on the fitnesses of individuals who are likely to actually exist in a real population. To do this we employ the single-site model of (5) and adopt the following assumptions to move from a single site to a whole-genome analysis: all functional sites share the same values for *u* and *s* described above; there is a multiplicative fitness relationship among functional sites; and there is no linkage disequilibrium between functional sites.

We first find the distribution of the whole-genome fitness *W* of a randomly chosen individual whose genome comprises 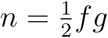 functional sites, when site fitness and frequency values are as given in (5). The fitness of an individual who is *AA* at *x* sites, *AM* at *y* sites and *MM* at *z* sites (*x* + *y* + *z* = *n*) is 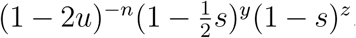. The probability that an individual is *AA* at *x* sites, *AM* at *y* sites and *MM* at *z* sites is

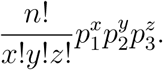

From this the mean of *W* is, from multinomial distribution formulae,

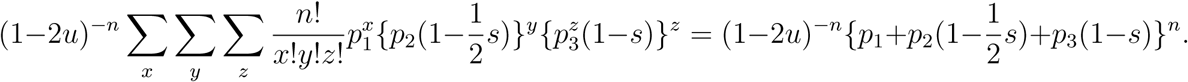

Given the values in (5), the mean of *W* is,

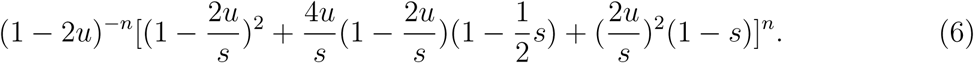

The variance of *W* is

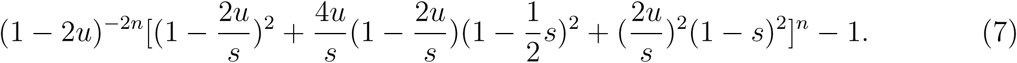

The mean of *W* is (1 − 2*u*)^−*n*^(1 − 2*u*)^*n*^ = 1, as expected. If terms of order *u*^2^ are ignored, the variance of *W* is

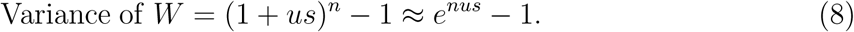

When parameter values are as given in (3) and *f* = 0.05 this variance is about 0.00803, so that the standard deviation in fitness is about 0.0896. The idealized fittest possible individual has fitness (1 − 2*u*)^−*n*^ ≈ 4.95, about 44 standard deviations above the mean. This corresponds to the previously calculated probability *e*^−320^ that such an individual exists. The same conclusion arises for other plausible parameter values. No such idealized individual will arise in practice.

The fitness 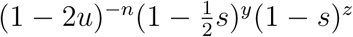 referred to above does not have a normal distribution. However the logarithm of this fitness, namely 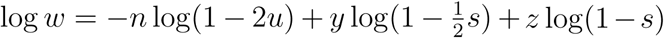 can be taken as having a normal distribution since for all practical purposes both *y* and *z* approximately have a normal distribution. (All logarithms are natural.) Therefore, to a sufficiently accurate approximation, the fitness *W* of an individual taken at random has a lognormal distribution (as illustrated in Figure 1, most clearly for the case *s* = 0.01).

**Figure 1:**
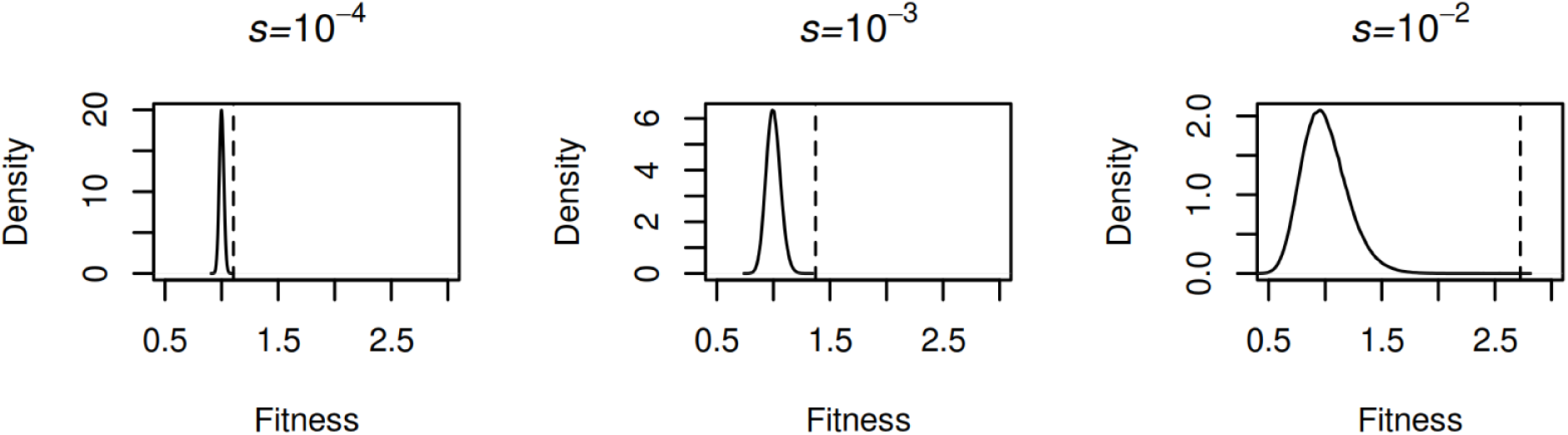
Simulated fitness distributions for the parameters *u* = 5 × 10^−9^, *g* = 6.4 × 10^9^, and *f* = 0.25. The dashed line shows the fitness of the fittest individual likely to exist in a population of size *N* = 10^9^, as given by (12).

The mean *µ* and the variance *σ*^2^ of log *W* can be found from the known mean 1 of *W* and known variance of *W* given in (8). These give

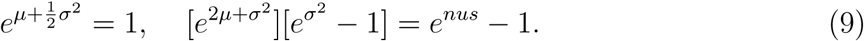

From this,

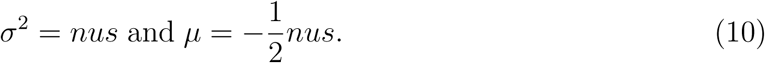

The above calculation can be confirmed by that of Lesecque *et al.* (2012), who use different notation and a slightly different variance formula. Lesecque *et al.* consider the case *nu* = 10 (equivalent, for the parameters given in (3), to *f* = 0.625), 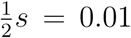 and calculate the probability that *W* takes a value between 0.5 and 2 to be 97%. This is the probability that log *W* takes a value between − log 2 = −.69315 and and + log 2 = +.69315. With the values *nu* = 10 and 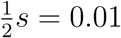, log *W* has mean −0.05 and variance 0.1. The probability that log *W* takes a value between − log 2 and and + log 2 is found from the normal distribution to be 97%, in agreement with the calculation made by Lesecque *et al.* (2012). Thus only about 3% of individuals have a fitness outside the range 0.5 to 2.0 for these parameter values. Calculations using other plausible parameter values lead to the same conclusion, namely that the variance of the fitness *W* is small and that the great majority of individuals have an easily achieved fitness.

#### 3.1.2 The fitness of the fittest individual likely to exist

We now calculate, for the additive case, the fitness of the fittest individual who is likely to exist in a reproducing population of a plausible size. The mean number 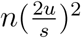 of *MM* sites in a randomly chosen individual is of order *u*^2^ and thus is very small, so that *MM* sites can be ignored in the calculations. Lesecque *et al.* (2012) also do this. The focus is therefore on *AA* and *AM* sites.

If terms of order *u*^2^ are ignored, the mean number of *AA* functional sites in a randomly-chosen individual is 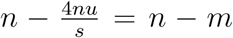 and the mean number of *AM* functional sites in a randomly chosen individual is *m*, where 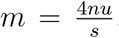. We assume that the fittest individual likely to appear in the population is *AA* at *n* − *m* + *r* functional sites and is *AM* at *m* − *r* functional sites, where the value of *r* has to be determined. To calculate *r* we assume that the number of functional sites at which a randomly chosen individual is *AM* has a Poisson distribution with mean *m*. (Lesecque *et al.* (2012) also make this Poisson distribution assumption.) This assumption implies that the standard deviation of this number of sites is 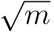. We conservatively assume a population of reproducing individuals of size 10^9^. Using the normal approximation to the Poisson, the fittest individual who is likely to appear in a reproducing population of size 10^9^ will therefore be *AA* at about 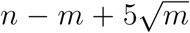 sites and *AM* at about 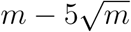 sites. Thus 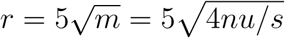.

It follows that when the mean population fitness is 1 the fitness requirement for the fittest individual who is likely to appear in a population of size 10^9^ is

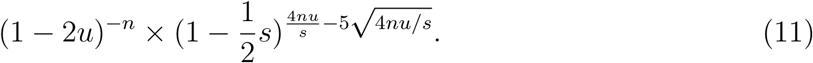

Since (1 − 2*u*)^−*n*^ ≈ *e*^2*nu*^, 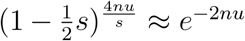 and 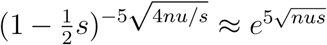, this is approximately

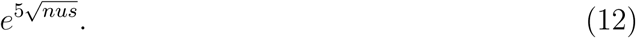

This is the fitness requirement for the value of *w*_*max*_ that is most likely to be empirically relevant to the human population, under our model. Figure 1 illustrates this value for *f* = 0.25 and varying *s*, and Table 1 presents the value of *w*_*max*_ likely to actually occur according to (12) for a range of values for *f* and *s*. For both Figure 1 and Table 1, *g, u*, and *N* have the values given in (3). The strong influence of *s* on the result is apparent: only for the highest value of *s* and the highest values of *f* does there seem to be any potential difficulty in realizing the values of *w*_*max*_ presented in Table 1. Thus we conclude that, first, there is not any very strong case for limiting the functional fraction from a mutational load standpoint; and, second, any such argument depends strongly on, and must take into account, the selection coefficients of newly arising deleterious mutations.

### 3.2 Comparison of results

We now compare the above findings to those of Graur (2017). The fertility requirements computed by Graur (2017) for the additive case are implicitly based on the fitness of an idealized individual. As we show above, an individual who is *AA* at all sites is vanishingly unlikely to exist.

The values in (5) show that the maximum fitness possible, that of an “optimal” individual who is *AA* at all *n* functional sites in the genome, is *w*_max_ = (1 − 2*u*)^−*n*^ ≈ *e*^2*nu*^. Since the mean fitness is 1, this value is a measure of the quantity defined in (2)

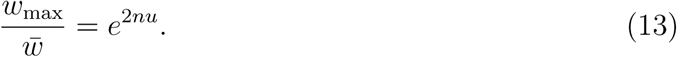

We confirm this calculation is essentially the same made by Graur (2017) for the case *f* = 0.10. Graur assumes that *g* = 6.114 × 10^9^ so that according to our model 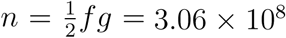. When *u* = 5 × 10^−9^, *e*^2*nu*^ = *e*^3.057^ = 21. For the case *f* = 0.20, with the other parameter values unchanged, *e*^2*nu*^ = 452. These values (21 and 452) are nearly the same as the tabled values in Graur (2017) for *f* = 0.05 and *f* = 0.10, respectively. The factor-of-two difference in *f* is due to the fact that Graur, erroneously in our view, treats the total number of sites as the number of haploid sites, not the number of diploid sites, *i.e.* omits the factor 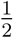 in 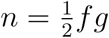.

With the numerical values in (3) and with *f* = 0.05 the expression in (13) is about 4.95, substantially higher than the 1.56 that results from the use of (12).

Graur argues that the quantity defined in (13) cannot be higher than some reasonable value for humans. He interprets this quantity as the mean fertility, that is, the average number of offspring per adult conditional upon survival to reproduction. He sets this maximum value at 1.8, based on historical data, which corresponds to 3.6 offspring per mating pair. This limits 2*nu* to about 0.6. Since 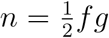 this sets a limit on the value of *f*. With the values *g* and *u* given in (3) this limiting value for *f* is about 0.02—quite low indeed.

Our interpretation of the quantity 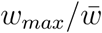 is more liberal than Graur’s: we do not interpret it as mean requisite fertility, because we are not using a pure viability selection model, but as the fitness of the fittest individual. Thus our interpretation of the approach of Graur yields somewhat higher possible values for *f* than occur in Graur (2017), as shown in the bottom row of Table 1, but still almost certainly no higher than *f* = 0.10 given the parameter values in (3).

### 3.3 The recessive case

We next discuss the recessive case, in which the single-site fitnesses differ from (4) in that the fitness of the heterozygote is equal to the fitness of the *AA* homozygote. As in the additive case, we normalize so that the mean fitness is 1. This leads to the following fitness table.

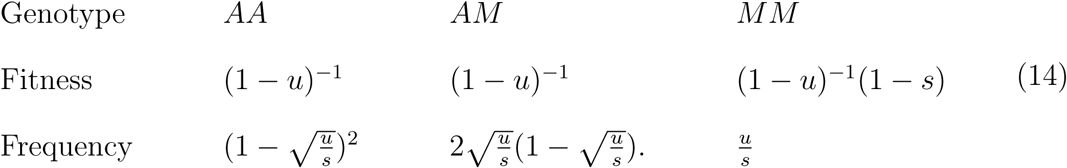

As in the additive case, an individual of the highest possible theoretical fitness will never exist in any population of a size relevant to humans. The probability that an individual taken at random from the population is either *AA* or *AM* at all *n* functional sites in the genome is 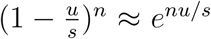. If we assume the parameter values in (3) and put *f* = 0.05, this probability is *e*^−80^. The same conclusion is reached with other reasonable parameter value choices.

We next calculate realistic whole-genome fitnesses making the same simplifying assumptions as those made for the additive case.

#### 3.3.1 The distribution of whole-genome fitness

We define *W* as the fitness of an individual taken at random. With site fitnesses as given in (14) and the various assumptions made above, the whole-genome mean of *W* is 1 and the the whole-genome variance of *W* is found from (14) and multinomial distribution formulae to be

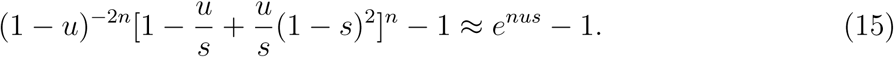

This leads to the same asymptotic formula (8) that applied in the additive case. The fitness of an individual who is either *AA* or *AM* at all functional sites is (1 − *u*)^−*n*^. When *f* = 0.05 and other parameter values are as in (3), this is about *e*^0.8^ ≈ 2.23, ∼14 standard deviations above the mean. As stated for the additive case, such an individual will never exist. The same conclusion holds for other plausible parameter values.

The fitness *W* of a randomly-chosen individual who is *AA* at *x* sites, *AM* at *y* sites, and *MM* at *z* sites is (1 − *u*)^−*n*^(1 − *s*)^*z*^. Thus the fitness of this individual does not have a normal distribution. However to a close approximation log *W* = −*n* log(1 − *u*) + *z* log(1 − *s*) has a normal distribution since *z* has approximately a normal distribution. Therefore, to a close approximation, *W* has a lognormal distribution. The mean *µ* and variance *σ*^2^ of log *W* can be found from the known mean (1) and known variance (*e*^*nus*^ − 1) of *W* using standard formulas relating parameters in a normal distribution and the parameters in the corresponding lognormal distribution. It is found that

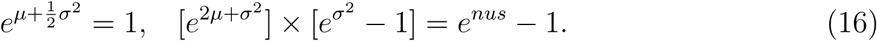

From this, *σ*^2^ = *nus* and then 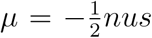. These are the same formulae as found for the additive case in (10).

#### 3.3.2 The fitness of the fittest individual likely to exist

We now find the fitness of the fittest individual likely to appear in the population. Since there are *n* functional sites and the probability that an individual is *MM* at a given functional site is *u/s*, the mean number of *MM* sites in a randomly chosen individual is *k* = *nu/s*. We assume that the actual number of *MM* sites carried by a randomly chosen individual has a Poisson distribution with parameter *k*. The standard deviation of this distribution is 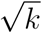. This distribution can be approximated by a normal distribution with mean *k* and standard deviation 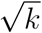. The properties of the statistics of extreme values of normal random variables show that in a population of 10^9^ reproducing individuals, the individual with the smallest number of *MM* sites will have about 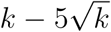 such sites, or 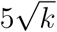 fewer than the mean (of *k*). It follows that this individual is *AA* or *AM* at 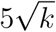 sites more than the mean number *n* − *k* of these sites. From (14) the fitness of an individual having this number of *AA* or *AM* sites is

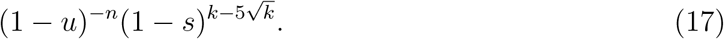

This is approximately 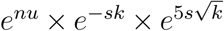. Since 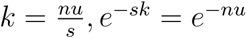 and 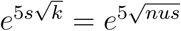, the expression in (17) becomes

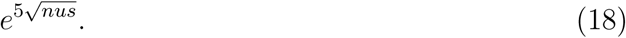

Note that this is exactly the same formula as in the additive case (12), so that the results shown in the first three rows of Table 1 apply to the recessive case as well as the additive case. Similarly, Figure 1 is illustrative for both cases.

## 4 Discussion and further considerations

### 4.1 The Wright-Mayr viewpoint

A main point of this paper is that no individual with the theoretical maximum fitness, given the fitness model, will ever exist in a real population. This point is not new. It was made by Wright (1977, page 481) in his discussion of Haldane’s (1957) evolutionary, or substitutional, load concept, which was also based on a non-existent “optimal” individual. Wright states that “if many loci are involved, the genotype that combines the [optimal] genotypes at all loci is in general so rare theoretically that neither it nor anything approaching it exists in a finite population.” Dobzhansky (1957), in response to Muller (1950), noted that for both flies and humans, “perhaps [individuals with no deleterious mutations] would be a superfly and a superman, but the fact is that such have never existed on earth.” Mayr (1970) makes the same point in stating that “the whole approach [to Haldane-based load calculations] is misleading. It is based on a set of assumptions that have no real validity, primarily that of the existence of an optimal homozygous genotype.” Lesecque *et al.* (2012) describe an individual of optimal homozygous genotype as an “idealized” individual and Agrawal and Whitlock (2012) state that such an individual is unlikely to exist. Charlesworth (2013) states that “the mean fitness of a population relative to the fitness of a hypothetical optimal genotype that has a very low chance of being present in the population is essentially irrelevant.” Henn *et al.* (2015) describe calculations involving *w*_max_ as idealized, and refer to the challenge of finding an empirically relevant *w*_*max*_. Our equation (12) is relevant to this question.

The Wright-Mayr view, and that of the authors cited above, is the one adopted here. Wright also noted that Crow’s (1970) definition of load (equation 1) is flexible in that it relates to the”fitness requirement of actually or theoretically available [geno]types”. At the whole-genome level, “actually available” concerns real populations and “theoretically available” concerns idealized populations. We believe that the appropriate choice is “actually available,” which is the Wright-Mayr viewpoint.

### 4.2 Remarks on the Haldane load

Agrawal and Whitlock (2012) define the load as in (1), so that for them the load in the additive case is *L* = 1 − *e*^−2*nu*^, and make two comments about this formula. First, they state that the fact that this load formula is independent of *s* has led to a “misleading sentiment” among theoretical population geneticists who then feel that nothing need be known about the value of *s* or about ecological considerations in assessing loads. We agree, and believe that load calculations that ignore *s* are not realistic (see Table 1) and have influenced population genetics theory for far too long. Second, they state that there is very little empirical evidence that Haldane’s (1957) load theory, based on the formula *L* = 1 − *e*^−2*nu*^, is even approximately correct. A likely reason for this lack of evidence is that Haldane’s theory, being based on this formula, is not empirically relevant to populations with parameters similar to those for humans.

In humans, *nu* is on the order of 1 to 10. For most microbes, in contrast, *nu* ≪ 1. For example, for *Escherichia coli*, estimates of *nu* are on the order of 10^−4^ (Kibota and Lynch 1996). If *s* = 0.001, then ∼90% of such a bacterial population would have no deleterious mutations, in contrast to the case of human populations in which no individual with zero deleterious mutations would ever occur. For many microbes, then, Haldane load theory may be appropriate.

### 4.3 The stochastic effect of finite population size

In the calculations above we have in effect assumed in the recessive case that the mean number of *MM* sites in a randomly chosen individual is 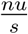. This calculation ignores stochastic effects in a population of finite size. Lesecque *et al.* (2012) show, for the recessive case, that when these effects are taken into account a slightly more accurate expression for this mean is 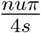. This adjustment would not materially change the main points of our analysis, and we suspect that a similar result holds for the additive case. We caution that any stochastic model (the Wright-Fisher model or alternatives) must make assumptions that are unlikely to be accurate for a real population, so that any inferences into the differences between load in finite and infinite populations are of limited value.

### 4.4 The value of *N*

Our choice *N* = 10^9^ is intended to be extremely conservative. Of all the parameters involved, the value chosen for *N* in a population that has subdivided and increased substantially in size over hundreds of thousands of years is possibly the most problematic. All the calculations above, and many in the literature, assume random mating. Henn *et al.* (2015) consider in detail the fact that for many thousands of years random mating is an unreasonable assumption, given the division of the human population into different sub-groups, based largely on geographical dispersion. This dispersion also bears on the reasonable choice for *N*. Henn *et al.* also consider complications due to the effects of population-size bottlenecks and the “prodigious rate” of growth in the size of the human population, increasing from a few hundred thousand about 13,000 years ago to several millions four thousand years ago. The conclusions that we reach about possible values of *f* continue to hold, and are strengthened by, any reasonable choice for the various values of the human population size over the last two hundred thousand years.

### 4.5 Other fitness models

When *h* is positive a fitness model generalizing (5) is

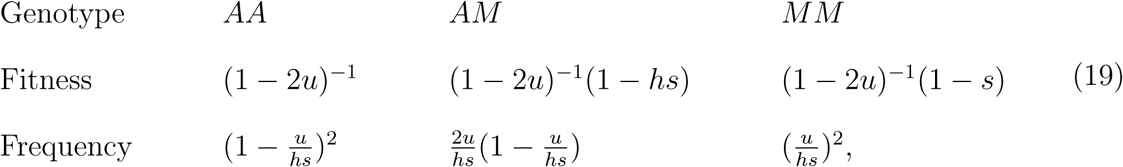

The case 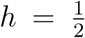 corresponds to the additive model. The “Haldane load” (1 − 2*u*)^−*n*^ is independent of *h*. It might then be expected that the realistic load generalizing (12) is also independent of the value of *h*, but this is not so. It is found after some algebra that with fitness and frequency values as given in (19), the mean of *w* is 1 but the the variance of *w* is no longer as given in (8) but is, instead, *e*^2*nuhs*^ − 1. From this the mean *µ* and variance *σ*^2^ of log *w* are no longer as given in (8) but are, instead,

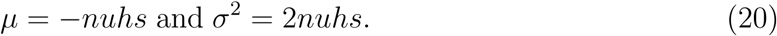

Henn *et al.* (2015) state that the average value of *h* in (19) across various non-human organisms is about 0.25. If the average value *h* = 0.25 applies also in humans, these amended values strengthen the conclusions reached in this paper for the additive case.

### 4.6 Stochasticity of number of offspring

Two individuals with the same intrinsic fitness do not necessarily have the same number of offspring: stochastic effects have to be taken into account. Lesecque *et al.* (2012) quantify this by discussing three models: an asexual model, a monogamous diploid model, and a freely interbreeding diploid model, and for each model calculate the probability *P*(0) that an individual has (in the monogamous diploid case, a couple have) no offspring. They consider the effect of the value of *nu* on *P*(0) for a given value of *s* and produce very interesting graphs describing this effect (their Figure 4). Our interest is the effect of *s* on *P*(0) for a given value of *nu*. The graphs in Figure 4 of Lesecque *et al.* (2012) show that the value of *P*(0) increases very slowly with *s* in all three cases. The number of offspring for an individual is determined more by stochastic effects than by the individual’s intrinsic fitness. The reason for this is the fact that the variance in fitness as given in (10) is very small.

### 4.7 Other kinds of loads

Load-based arguments seeking to limit the value of *f* need not remain limited to the mutational load. The substitutional load and the segregational loads also depend to some extent upon *f* and might be considered as well. The criticism of load arguments by Wright, Mayr and others referred to above were made with respect to one or both of these loads. If these loads were taken into account as well as the mutational load the possible values of *f* would be smaller. However these load calculations are subject to the same criticisms that we have made for the mutational load.

### 4.8 Generalization to *s̄*

In this section we extend our analysis to consider several classes of deleterious mutations. Suppose that there are *k* different mutant types *M*_1_, *M*_2_, *…, M*_*k*_, having respective mutation rates *u*_1_, *u*_2_, *…, u*_*k*_ and relative fitnesses 1 (for *AA*), 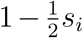 (for *AM*_*i*_) and 1 − *s*_*i*_ (for *M*_*i*_*M*_*i*_). We assume that each of these mutants is in mutation-selection balance, so that the frequency of *AA* is 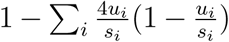, the frequency of *AM*_*i*_ is 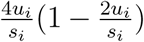 and the frequency of *M*_*i*_*M*_*i*_ is 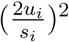. Fitnesses are now normalized so that the mean fitness is 1. This leads to the fitness values (1 − 2*u*)^−1^ for *AA*, 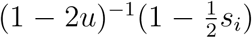 for *AM*_*i*_, and (1 − 2*u*)^−1^(1 − *s*_*i*_) for *M*_*i*_*M*_*i*_, where *u* = ∑*u*_*i*_.

Under the assumptions made in the paper the whole-genome variance in fitness is then

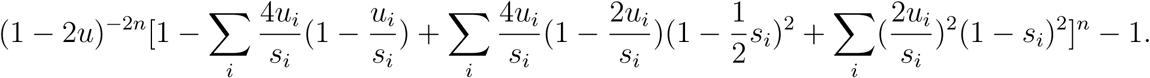

If terms of order 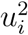 are ignored, this variance is

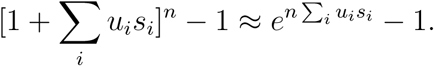

This is a generalization of Equation (8). It follows that where *nus* occurs above the more general expression *n* ∑_*i*_ *u*_*i*_*s*_*i*_ can be written. Similarly the expression *s* appearing in Table 1 can be written more generally as 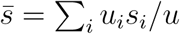.

The fitness *W* of an individual taken at random does not have a normal distribution. However to a close approximation log *W* has a normal distribution. The mean *µ* and variance *σ*^2^ of this distribution can be found immediately from (9) by replacing *nus* by 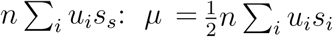 and *σ*^2^ = *n* ∑_*i*_ *u*_*i*_*s*_*i*_.

## 5 Summary and Conclusions

The per-nucleotide rate of mutation and the total size of the human genome appear to be fairly well established. The fraction *f* of the human genome that is functional remains uncertain. We have shown that when considering the likely maximum realized fitness in a finite population, the limit to *f* is by no means low. This result stands in contrast to arguments that depend upon the fitness of an individual who possesses the theoretical maximum fitness of the particular model employed. Such arguments appear to establish a rather low limit for *f*, but suffer from the flaw that such an individual is only vanishingly likely to exist. Calculations that purport to establish a load should, in our view, be based on the distribution of actual fitnesses that are expected to exist in a real population. As we have shown, the properties of this distribution depend not just on *nu* (the number of *de novo* deleterious mutations per individual) but also on *s*, the selection coefficient against deleterious mutations.

Using the approach of Graur (2017) and adopting the most plausible value for the human per-base-pair deleterious mutation rate, the limit to *f* is ∼2% − 10%. In contrast, we have shown that when considering the likely maximum realized fitness in a finite, persisting human population, much higher values for *f*, with considerable uncertainty introduced by the unknown value of the parameter *s*, are plausible (Table 1).

We stress that we, in this work, take no position on the actual proportion of the human genome that is likely to be functional. It may indeed be quite low, as the contemporary evidence from species divergence and intraspecies polymorphism data suggests. Many of the criticisms of the ENCODE claim of 80% functionality (e.g. Doolittle 2013, Graur 2013) strike us as well-founded. Our conclusion is simply that an argument from mutational load does not appear to be particularly limiting on *f*.

## Literature cited

Agrawal, A., Whitlock M. 2012. Mutation load: the fitness of individuals in populations where deleterious alleles are abundant. Ann. Rev. Ecol. Evol. 43: 115–135.

Besenbacher, S. et al. 2015. Novel variation and *de novo* mutation rates in population-wide de novo assembled Danish trios. Nature Communications 6: 5969.

Boyko, A. et al. 2008. Assessing the evolutionary impact of amino acid mutations in the human genome. PLoS Genet. 4: e1000083.

Charlesworth, B. 2013. Why are we not dead one hundred times over? Evolution 67(11): 3354–3361.

Crow, J. 1970. Genetic loads and the cost of natural selection. In Mathematical topics in population genetics, ed. K. Kojima. Springer-Verlag.

Crow, J. F., & Kimura, M. 1970. An Introduction to Population Genetics Theory. Harper & Row.

Dobzhansky, T. 1957. Genetic loads in natural populations. Science 126: 191–194.

Doolittle, W. F. 2013. Is junk DNA bunk? A critique of ENCODE. PNAS 110: 5294–5300.

Eyre-Walker, A., Woolfit, M., Phelps, T. 2006. The distribution of fitness effects of new deleterious amino acid mutations in humans. Genetics 173: 891–900.

Eyre-Walker, A., Keightley, P. 2007. The distribution of fitness effects of new mutations. Nature Reviews 8: 610–619.

ENCODE Project consortium 2012. An integrated encyclopedia of DNA elements in the human genome. Nature 489: 57–74.

Graur, D. et al. 2013. On the immortality of television sets: “function” in the human genome according to the evolution-free gospel of ENCODE. Genome Biology and Evolution 5(3): 578–590.

Graur, D. 2017. An upper limit on the functional fraction of the human genome. Genome Biology and Evolution 9: 1880–1885.

Gregory, T. R. 2005. Synergy between sequence and size in large-scale genomics. Nature Reviews Genetics 6(9): 699–708.

Gulko, B. et al 2015. A method for calculating probabilities of fitness consequences for point mutations across the human genome. Nature genetics 47(3): 276.

International Human Genome Sequencing Consortium 2001. Initial sequencing and analysis of the human genome. Nature 409(6822): 860–921.

Kibota, T. and Lynch, M. 1996. Estimate of the genomic mutation rate deleterious to overall fitness in *E. coli*. Nature 381(6584): 694–696.

de Koning, A. J. et al 2011. Repetitive elements may comprise over two-thirds of the human genome. PLOS Genetics 7(12), e1002384.

Haldane, J. 1937. The effect of variation on fitness. Amer. Nat. 71: 337–349.

Haldane, J. 1957. The cost of natural selection. J. Genet. 55:511–524.

Henn, B. et al. 2015. Estimating the mutation load in human genomes. Nature Reviews Genetics 16(6): 333–343.

Lesecque Y., Keightley P., Eyre-Walker A. 2012. A resolution of the mutation load in humans. Genetics 191: 1321–1330.

Mayr, E. 1970. Populations, Species and Evolution. Harvard University Press.

Meader, S. et al 2010. Massive turnover of functional sequence in human and other mammalian genomes. Genome research 20(10): 1335–1343.

Milholland B. et al. 2017. Differences between germline and somatic mutation rates in humans and mice. Nature Communications 8(1): 1–8.

Muller, H. J. 1950. Our load of mutations. Am J Hum Genet. 2: 111–176.

Ponting, C. and Hardison, R. 2011. What fraction of the human genome is functional? Genome research 21: 1769–1776.

Rands, C. et al. 2014. 8.2% of the human genome is constrained: variation in rates of turnover across functional element classes in the human lineage. PLOS Genetics 10(7), e1004525.

Schneider V. et al. 2017. Evaluation of GRCh38 and de novo haploid genome assemblies demonstrates the enduring quality of the reference assembly. Genome Research 27.5: 849–864.

Thomas, C.A. Jr. 1971. The genetic organization of chromosomes. Annu. Rev. Genet. 5: 237–256.

Ward, L.D. and Kellis, M., 2012. Evidence of abundant purifying selection in humans for recently acquired regulatory functions. Science 337:1675–1678.

Wright, S. 1977. Evolution and the Genetics of Populations 3. Experimental results and evolutionary deductions. University of Chicago Press.

